# Characterizing the interaction of a type VII-secreted antimycobacterial toxin with its small helical partner proteins

**DOI:** 10.64898/2026.08.24.746431

**Authors:** Eunice K.E. Lee, Kieran Bowran, Eleanor R. Boardman, Tracy Palmer

## Abstract

The type VII secretion system (T7SS) is a membrane-embedded protein export pathway found in mycobacteria and Gram-positive bacteria. Recently it was shown that *Mycobacterium abscessus* uses its ESX-4 variant of the T7SS to secrete a toxin, EatA, which targets arabinogalactan present in the mycobacterial cell envelope. Prior to its export, EatA forms a complex with a pair of small proteins from the WXG100 family, TapA1 and TapA2. Here we investigated a structural model of the EatA N-terminal domain in complex with TapA1 and TapA2 using site-directed mutagenesis and bacterial 2-hybrid assays. Our results are consistent with the three proteins forming a stacked bundle of α-helices. Structural modelling also predicted an interaction of the EatA-TapA1-TapA2 complex with EsxT-EsxU, a second pair of WXG100-family proteins that are likely required for the mechanistic operation of ESX-4. Whilst we could demonstrate a potential interaction between TapA2 and EsxT by bacterial 2-hybrid analysis, we were not able to purify a complex of all five proteins.

## Introduction

Bacterial protein secretion systems play essential roles in nutrient acquisition, colonisation and virulence. Gram-negative bacteria encode a plethora of different secretion systems that mediate transport across the double-membraned cell envelope (1). However, these Gram-negative systems are generally not found among the Gram-positive bacteria or diderm mycobacteria, which instead have their own specialised system, termed the type VII secretion system (T7SS) (2-6).

The composition of the T7SS is highly variable across bacterial species and has been divided into ten subtypes, T7SSa – T7SSj (7). The mycobacterial system is classified as the T7SSa, also termed ESX (for ESAT-6 secretion system), and is the best-studied of the ten systems. Up to five distinct T7SSa systems, ESX-1 - ESX-5 can be found in a single mycobacterial species. *Mycobacterium tuberculosis* has all five ESX systems, with ESX-1, ESX-3 and ESX-5 being closely linked with virulence (8). Fast growing mycobacterial species such as *Mycobacterium smegmatis* lack ESX-2 and ESX-5, while *Mycobacterium abscessus* possesses just the ESX-3 and ESX-4 systems (9).

The ESX secretion machinery is a 2.3 MDa complex situated in the cytoplasmic membrane (10). At the centre of the system is a hexameric arrangement of the ATPase EccC. Each EccC monomer has two transmembrane domains at its N-terminus, followed by four cytosolic FtsK/SpoIIIE-like ATPase domains. The EccC hexamer forms the secretion pore, and likely powers protein secretion through ATP hydrolysis (10-12). EccC also has a key role in substrate recognition through its most distal ATPase domain (3, 13, 14).

The activity of ESX systems depends on a canonical pair of proteins from the WXG100 family that are usually encoded at the same genetic locus as the secretion machinery (e.g. 3, 15-18). WXG100 proteins are approximately 100 amino acids in length and adopt a helical hairpin fold, with the conserved WxG motif located at the hairpin turn (19). In the ESX-1 system these proteins are EsxA and EsxB, and they form an antiparallel heterodimer that is secreted by ESX-1 (2-4, 20). Deletion of *esxBA* abolishes ESX- 1-dependent secretion (e.g. 18), indicating that the proteins are mechanistically essential for operation of the secretion system. The mechanistic requirement for a WXG100 dimer has also been shown in the T7SSb system (21). In this case a single WXG100 protein, EsxA forms an antiparallel homodimer, and is required for export of other known substrates (21, 22).

In addition to WXG100 proteins that are specifically linked with the T7SS secretion machinery, other WXG100-like proteins are also encoded in bacterial genomes. These are usually found adjacent to genes that encode larger substrates of the T7SS, often antibacterial toxins, and have been described in the T7SSb system of *Staphylococcus aureus* and *Streptococcus intermedius* (e.g. (23-27)). Unlike EsxA, these WXG100 family proteins are specific for the toxin with which they are co-encoded and they bind to its helical trafficking domain to generate a secretion-competent complex (26-28). The WXG100- like partners may be subsequently secreted along with their toxin partner, most likely as a complex (27-29).

It has recently been reported that antibacterial toxins can also be substrates of the T7SSa (30). The ESX-4 system of *M. abscessus* was shown to secrete an arabinanase toxin, EatA, that degrades the arabinogalactan component uniquely present in the cell envelopes of Corynebacteriales. To protect from self-intoxication, *M. abscessus* strains also produce EatI, an immunity lipoprotein that binds tightly to EatA to block its activity. EatA is encoded at a genetic locus with two small genes for WXG100-like proteins, named TapA1and TapA2 (for **t**oxin **<u>a</u>**ccessory **<u>p</u>**rotein) that bind to the N-terminal domain of EatA (30). Here we investigate the interaction of TapA1 and TapA2 with EatA through a programme of site-directed mutagenesis.

## Methods

### Bacterial Strains and growth conditions

The following *E. coli* strains were used in this work: DH5α (F^-^ φ80*lac*ZΔM15 Δ(*lac*ZYA-*arg*F)U169 *rec*A1 *end*A1 *hsd*R17(r_K_^-^, m_K_^+^) *pho*A *sup*E44 λ^-^*thi*- 1 *gyr*A96 *rel*A1; Promega) for general cloning, BTH101 (F^-^ *cya-99 araD139 galE15 galK16 rpsL1 (*Str^R^*) hsdR2 mcrA1 mcrB1*; (31) for bacterial 2-hybrid analysis, and M15 (pREP4) (F^-^ *lac ara gal mtl* [*kan*R, *lacI*]; Qiagen) for expression of proteins from the pQE70 vector. *E. coli* was grown in lysogeny broth (LB; per litre, 10g casein digest peptone, 10g NaCl, 5g yeast extract pH 7.0) or on LB agar at 37°C unless otherwise stated. Media were supplemented with ampicillin (Amp; 100 μg/mL) or chloramphenicol (Cml; 25 μg/mL) where necessary for plasmid maintenance.

### Plasmid construction

All plasmids used and constructed in this work are shown in Table 1. Plasmids pUT18-*tapA1*, pUT18-*tapA2*, pUT18-*eatA_NT_*, pT25-*tapA1*, pT25-*tapA2*, pT25-*eatA* and pT25-*eatA_NT_* have all been described previously (30). Gibson Assembly (either using NEBuilder HiFi DNA Assembly Master Mix (NEB, #E2621; 2X) or ClonExpress II One Step Cloning Kit (Vazyme, #C112-01)) was used to construct all plasmids in this work, with *M. abscessus* ATCC19977 chromosomal DNA as the template. Plasmid pQE70-*tapA1*-*tapA2*-*eatA* was constructed as a precursor to generate pQE70-*tapA1*- *tapA2-eatA-esxU-esxT*. Site-directed mutations were introduced into plasmids using Gibson Assembly. All oligonucleotides used in this work are listed in Table S1. Sanger sequencing was performed to confirm the correct inserts in plasmids either using Mix2Seq (Eurofins) or whole-plasmid sequencing (Plasmidsaurus).

**Table 1.** Plasmids used in this study.

| Plasmid | Description | Reference |
| --- | --- | --- |
| pUT18 | Bacterial 2-hybrid vector containing T18 *cya* gene fragment from *Bordetella pertussis*; Amp<sup>R</sup>. | (31) |
| pUT18-NarG | pUT18 producing NarG<sub>1-42</sub>-T18 fusion | (42) |
| pUT18-*tapA1* | pUT18 producing TapA1-T18 fusion | (30) |
| pUT18-*tapA1*<sub>V22D</sub> | pUT18 producing TapA1<sub>V22D</sub>-T18 fusion | This work |
| pUT18-*tapA1*<sub>I33D</sub> | pUT18 producing TapA1<sub>I33D</sub>-T18 fusion | This work |
| pUT18-*tapA1*<sub>L56D</sub> | pUT18 producing TapA1<sub>L56D</sub>-T18 fusion | This work |
| pUT18-*tapA2* | pUT18 producing TapA2-T18 fusion | (30) |
| pUT18-*tapA2*<sub>V47D</sub> | pUT18 producing TapA2<sub>V47D</sub>-T18 fusion | This work |
| pUT18-*tapA2*<sub>Y58D</sub> | pUT18 producing TapA2<sub>Y58D</sub>-T18 fusion | This work |
| pUT18-*tapA2*<sub>L69D</sub> | pUT18 producing TapA2<sub>L69D</sub>-T18 fusion | This work |
| pUT18-*eatA*<sub>NT</sub> | pUT18 producing EatA<sub>1-180</sub>-T18 fusion | (30) |
| pUT18-*eatA*<sub>NT(L29D)</sub> | pUT18 producing EatA<sub>1-180-L29D</sub>-T18 fusion | This work |
| pUT18-*eatA*<sub>NT(F53D)</sub> | pUT18 producing EatA<sub>1-180-F53D</sub>-T18 fusion | This work |
| pUT18-*eatA*<sub>NT(L121D)</sub> | pUT18 producing EatA<sub>1-180-L121D</sub>-T18 fusion | This work |
| pUT18-*eatA*<sub>NT(I128D)</sub> | pUT18 producing EatA<sub>1-180-I128D</sub>-T18 fusion | This work |
| pUT18-*eatA*<sub>NT(I136D)</sub> | pUT18 producing EatA<sub>1-180-I136D</sub>-T18 fusion | This work |
| pUT18-*eatA*<sub>NT(L155D)</sub> | pUT18 producing EatA<sub>1-180-L155D</sub>-T18 fusion | This work |
| pUT18-*esxT* | pUT18 producing EsxT-T18 fusion | This work |
| pUT18-*esxU* | pUT18 producing EsxU-T18 fusion | This work |
| pT25 | Bacterial two-hybrid vector containing T25 *cya* gene fragment from *B. pertussis*; Cml<sup>R</sup>. | (31) |
| pT25-NarJ | pT25 producing T25-NarJ fusion | (42) |
| pT25-*tapA1* | pT25 producing T25-TapA1 fusion | (30) |
| pT25-*tapA1*<sub>V22D</sub> | pT25 producing T25-TapA1<sub>V22D</sub> fusion | This work |
| pT25-*tapA1*<sub>I33D</sub> | pT25 producing T25-TapA1<sub>I33D</sub> fusion | This work |
| pT25-*tapA1*<sub>I36D</sub> | pT25 producing T25-TapA1<sub>I36D</sub> fusion | This work |
| pT25-*tapA2* | pT25 producing T25-TapA2 fusion | (30) |
| pT25-*tapA2*<sub>V47D</sub> | pT25 producing T25-TapA2<sub>V47D</sub> fusion | This work |
| pT25-*tapA2*<sub>Y58D</sub> | pT25 producing T25-TapA2<sub>Y58D</sub> fusion | This work |
| pT25-*tapA2*<sub>L69D</sub> | pT25 producing T25-TapA2<sub>L69D</sub> fusion | This work |
| pT25-*eatA* | pT25 producing T25-EatA fusion | (30) |
| pT25-*eatA*<sub>NT</sub> | pT25 producing T25-EatA<sub>1-180</sub> fusion | (30) |
| pT25-*eatA*<sub>NT(L29D)</sub> | pT25 producing T25-EatA<sub>1-180-L29D</sub> fusion | This work |
| pT25-*eatA*<sub>NT(F53D)</sub> | pT25 producing T25-EatA<sub>1-180-F53D</sub> fusion | This work |
| pT25-*eatA*<sub>NT(L121D)</sub> | pT25 producing T25-EatA<sub>1-180-L121D</sub> fusion | This work |
| pT25-*eatA*<sub>NT(I128D)</sub> | pT25 producing T25-EatA<sub>1-180-I128D</sub> fusion | This work |
| pT25-*eatA*<sub>NT(I136D)</sub> | pT25 producing T25-EatA<sub>1-180-I136D</sub> fusion | This work |
| pT25-*eatA*<sub>NT(L155D)</sub> | pT25 producing T25-EatA<sub>1-180-L155D</sub> fusion | This work |
| pQE70 | Plasmid for overexpression of proteins in *E. coli* from a *T5* promoter; Amp<sup>R</sup>. | Qiagen |
| pQE70-*esxU-esxT* | pQE70 encoding EsxU with a His<sub>6</sub> tag at the C-terminus and EsxT with a twin strep-tag at the C-terminus | This work |
| pQE70-*tapA1-tapA2* | pQE70 encoding TapA1 with a His<sub>6</sub> tag at the C-terminus and TapA2 with a twin strep-tag at the C-terminus | This work |
| pQE70-*tapA1-tapA2-eatA-esxU-esxT* | pQE70 encoding TapA1, TapA2 with a twin strep-tag at the C-terminus, EatA with a His<sub>6</sub> tag at the C-terminus, EsxU and EsxT with a HA tag at the C-terminus | This work |

### Bacterial-two-hybrid assay

This was carried out essentially as described previously (31). Plasmids pUT18 or pT25 encoding the appropriate bait or prey fusion proteins were co-transformed into *E. coli* BTH101. For each pairing, three independent colonies were inoculated into 5 ml of LB containing Amp and Cml and cultured overnight with shaking at 30°C. The following day, cultures were diluted to an OD600 = 1, and 10 μl was spotted onto fresh MacConkey agar plates containing Amp, Cml and 1% (w/v) D-maltose. Plates were incubated at 30°C until colonies were visible and positive control colonies were visibly pink (∼48 hours) before being photographed using the camera on an Apple iPhone XS Max or Apple iPhone 15 Pro.

### Recombinant protein purification

Unless otherwise specified, overexpression of proteins encoded on pQE70 plasmids was carried out as follows: an overnight culture of *E. coli* M15 (pREP4) harbouring the pQE vector of interest was refreshed at 1:100 dilution in LB/or terrific broth (TB) medium with appropriate antibiotics. Cultures were incubated at 37°C with shaking at 200 rpm until an OD600 of 0.6 – 1.0 was reached, after which they were supplemented with 1 mM IPTG. Cultures were further incubated for either 4 hours at 37°C or overnight for 18 hours at 18°C. Cells were harvested by centrifugation, washed once in PBS, and resuspended in Buffer A [50 mM 4-(2-hydroxyethyl)-1- piperazineethanesulfonic acid (HEPES), pH 7.5, 150 mM NaCl, 25 mM imidazole] supplemented with cOmplete EDTA-free protease inhibitor cocktail (Roche; #11836170001) and lysed by sonication. Clarified lysates were passed through a 5 ml HisTrap FF column attached to an ÄKTA Start system (Cytiva), pre-equilibrated with Buffer A. Following washing with 10 column volumes of Buffer A, proteins were eluted by applying a linear gradient of Buffer A containing 500 mM imidazole over 30 ml. To isolate proteins with a twin strep-tag, a 1 ml StrepTrap XT column attached to an ÄKTA Start system (Cytiva) was pre-equilibrated in Buffer B (100 mM Tris HCl pH 8.0, 150 mM NaCl, 1 mM EDTA). Fractions of interest from nickel affinity purification were diluted threefold using Buffer B before loading onto the column. The column was washed with 10 column volumes of Buffer B and bound proteins were eluted in 1x BXT Buffer (1 M Tris-HCl, 1.5 M NaCl, 10 mM EDTA, 500 mM biotin, pH 8.0; IBA Lifesciences). For size exclusion chromatography, samples from previous purification steps were concentrated using a Cytiva Vivaspin MWCO protein concentrator and separated on a Superdex 75 10/300 GL column linked to an ÄKTA Pure system (Cytiva). SEC was performed in a buffer comprising 25 mM HEPES pH 7.5, 150 mM NaCl.

### SDS-PAGE and western blot analysis

SDS-PAGE gels were electrophoresed at 80 V for 10 minutes followed by 200 V for 35 minutes, and proteins were visualised using InstantBlue Coomassie stain (Abcam). For western blotting, proteins were transferred from unstained SDS-PAGE gels to nitrocellulose membranes using transfer buffer (25 mM Tris, 192 mM glycine, pH 8.3, 20% methanol) at 250 mA for 1.5 hours. Following transfer, nitrocellulose membranes were blocked with 5% (w/v) skimmed milk powder suspended in TBST (20 mM Tris-HCl, pH 7.5, 150 mM NaCl, 0.1% (w/v) Tween-20) for at least an hour on a platform rocker. The milk suspension was replaced with fresh 5% (w/v) milk suspension in TBST containing the primary antibody and incubated at room temperature for an hour or at 4°C overnight, with gentle rocking. Three 10-minute washing steps with TBST were carried out before the secondary HRP-conjugated antibody was applied in fresh 5% (w/v) milk suspension in TBST for at least an hour on a platform rocker at room temperature. Three further 10-minute washing steps with TBST were carried out before Clarity Western ECL Western blotting substrate (Bio-Rad; #1705060), prepared according to the manufacturer’s instructions, was applied to the blots and immediately exposed in a G-Box Chemi XX6. The following primary antibodies were used: mouse α- His at 1:3,000 dilution (Invitrogen; #MA1-21315), mouse α-Strep at 1:5,000 dilution (Qiagen; #34850) and mouse α-HA at 1:3,000 dilution (Thermo Fisher Scientific; #26183). The secondary antibody was goat α-mouse IgG HRP conjugate, used at 1:5,000 dilution (Bio-Rad; #170-6516).

### Computational modelling of protein structures

Protein sequences were submitted to the AlphaFold3 server (https://alphafoldserver.com) for structural prediction (32). Models were visualised and analysed using UCSF ChimeraX v1.8 (https://www.rbvi.ucsf.edu/chimerax) (33).

## Results

### Heterologously-produced EsxT and EsxU form heterodimers but TapA1 and TapA2 do not

In mycobacteria it has been reported that WXG100 proteins associated with specific secretion systems, such as the ESX-1 pair EsxA and EsxB, form heterodimers (20). The WXG100 pairing for ESX-4 are EsxT and EsxU. Previous studies have shown that *M. abscessus* EsxT/EsxU also form a stable, soluble heterodimer with 1:1 stoichiometry (34), and AlphaFold3 modelling predicts a heterodimeric arrangement with high confidence (Fig 1a). To confirm that we can detect EsxT/EsxU heterodimers we co-produced these proteins in *E. coli* from a strong expression vector, with a C-terminal His_6_ tag on EsxT and a C-terminal twin strep (ts) tag on EsxU. The two proteins co-purified following sequential HisTrap and StrepTrap column chromatography in an apparent 1:1 ratio, as judged by the intensity of Coomassie blue staining following SDS PAGE (Fig 1b). To determine the approximate size of the EsxT- His_6_-EsxU-ts heterodimer we undertook size exclusion chromatography. Fig 1c shows that the complex eluted as a single species, and was close to the expected mass for a 1:1 stoichiometry.

**Fig 1.**
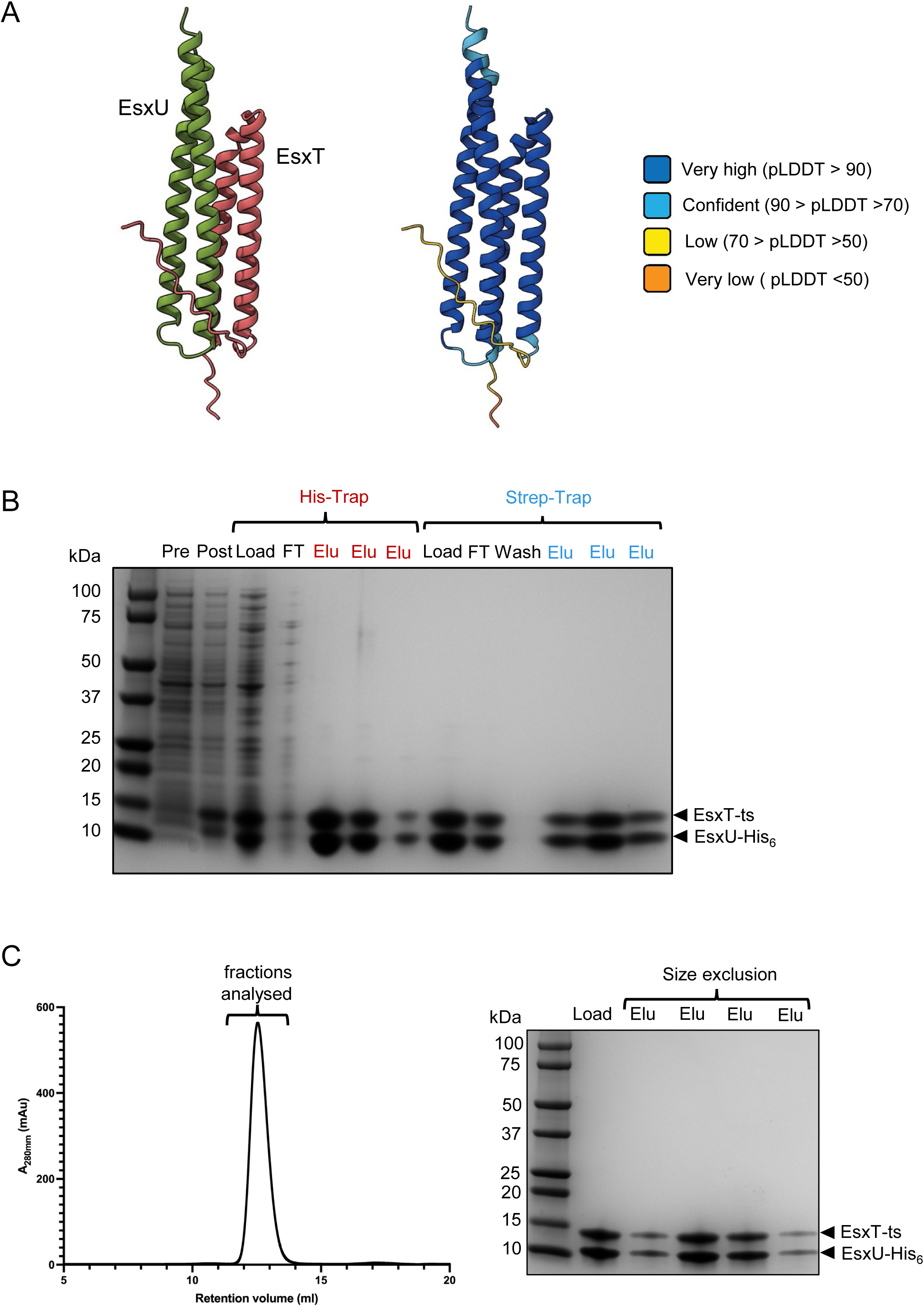
Co-purification of EsxT-ts and EsxU-His_6_. (a) AlphaFold3 model of the EsxT-EsxU complex coloured by chain (left) or pLDDT score (right). (b) SDS-PAGE analysis of EsxT-ts and EsxU-His_6_ purification. Lysates of *E. coli* co-producing EsxT-ts and EsxU-His_6_ purification were subjected to sequential purification by HisTrap and StrepTrap chromatography. Fractions were analysed by SDS- PAGE. Pre: pre-induction sample; Post: 4 hours post-induction sample; Load; sample loaded onto the column; FT: column flow-through; Wash: wash sample; Elu: elution samples from the peak fractions for each column. (c) *Left.* Size exclusion chromatogram of EsxT-ts and EsxU-His_6_ following StrepTrap and HisTrap purification. Comparison with the elution volume of protein standards gave a molecular weight estimate of 30 kDa for the complex. *Right*. SDS-PAGE analysis of peak fractions from size exclusion chromatography.

We next asked whether TapA1 and TapA2 could also form heterodimers. We noted that AlphaFold3 modelling predicted some degree of interaction, but with medium to low confidence (Fig 2a) and with a low predicted template modelling score (pTM = 0.32). Following a similar approach, we co-produced the two proteins in *E. coli* with a C-terminal His_6_ tag on TapA1 and a C-terminal ts tag on TapA2. After using the same culturing and expression conditions as those used for EsxT/EsxU, we prepared the soluble cell fraction and carried out HisTrap chromatography to isolate TapA1-His_6_ (Fig 2b). While abundant TapA1- His_6_ was purified by this method, we did not identify any TapA2-ts in our TapA1- His_6_ peak fraction, nor in the load sample, although some TapA2-ts was observed in the whole cell fraction post induction (Fig 2c). This would be consistent with TapA2-ts forming inclusion bodies, and indeed further analysis showed that we were able to detect TapA2-ts primarily in the insoluble fraction following cell lysis, which we could not improve even after varying cell growth media, or induction conditions (not shown). This finding suggests that TapA1 and TapA2 do not directly heterodimerise as they are found in different cell fractions, and is consistent with previous bacterial 2-hybrid analysis which does not support a direct interaction between TapA1 and TapA2 (30). These observations indicate that mycobacterial WXG100 family proteins can exhibit different behaviours.

**Fig 2.**
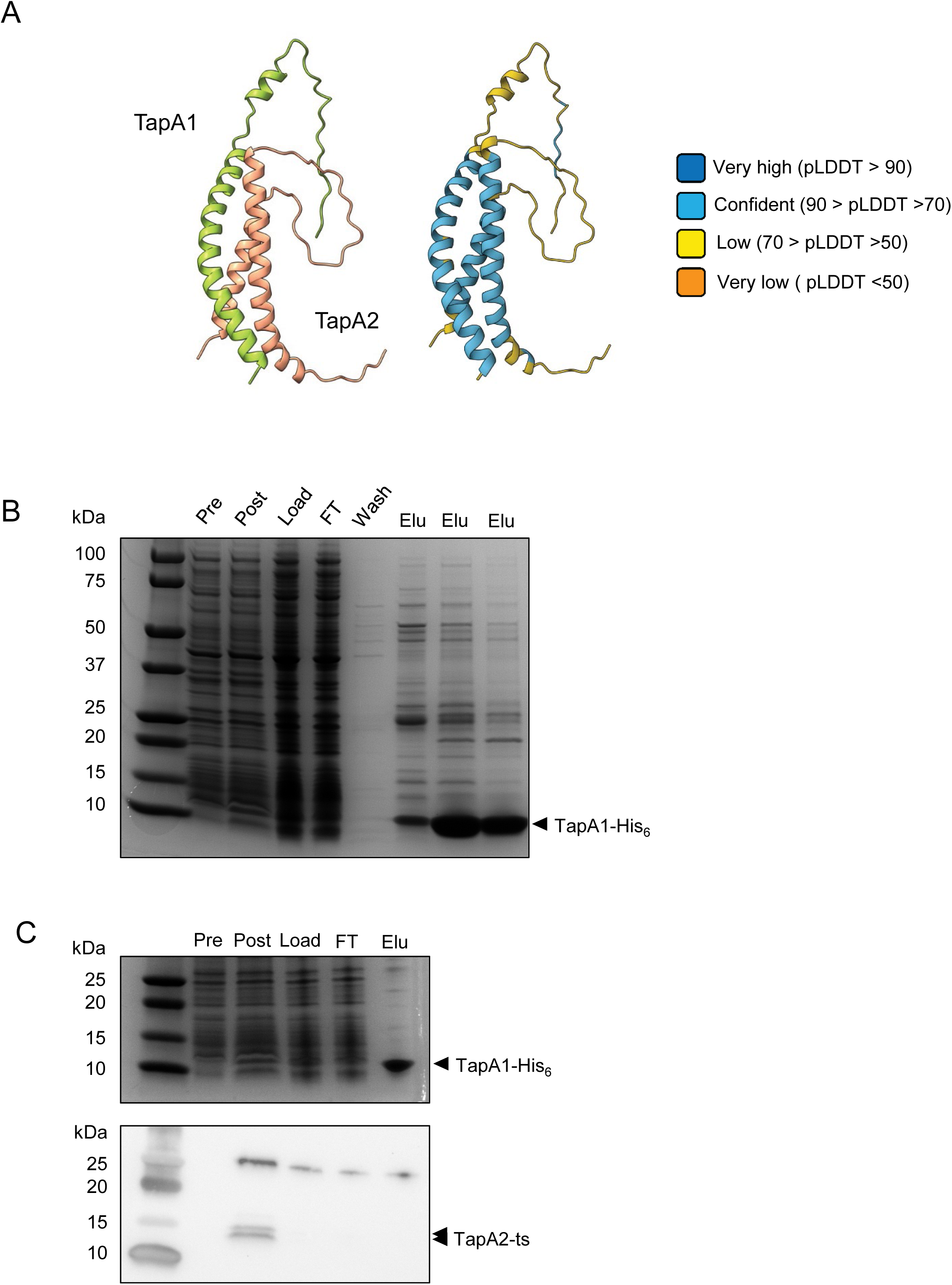
TapA1 and TapA2 do not co-purify. (a) AlphaFold3 model of a TapA1-TapA2 complex coloured by chain (left) or pLDDT score (right). (b) SDS-PAGE analysis of TapA1-His_6_ purification. Lysates of *E. coli* co-producing TapA1-His_6_ and TapA2-ts and were subjected to purification by HisTrap chromatography. Fractions were analysed by SDS-PAGE. Pre: pre-induction sample; Post: 4 hours post-induction sample; Load; sample loaded onto the column; FT: column flow-through; Wash: wash sample; Elu: elution samples from the peak fractions. (c) Western blot analysis using an α-Strep antibody to detect the presence of TapA2-ts in selected fractions from (b). The blot is shown in the bottom panel, and SDS-PAGE analysis of the same fractions is shown in the top panel.

### Experimentally testing an AlphaFold model of the TapA1-TapA2-EatA complex

In contrast to a poorly predicted TapA1/TapA2 heterodimer, AlphaFold3 modelling strongly predicts that these two proteins heterotrimerise with their co-encoded arabinanase toxin EatA (30). Co-purification and bacterial 2-hybrid studies have confirmed that the Tap proteins specifically interact with the helical N-terminus of EatA (EatA_NT_) ((30); Fig 3a). Within this model TapA1 and TapA2 do not make any direct contacts (Fig 3a). To probe the veracity of this model we undertook site-directed mutagenesis on residues predicted to form the interaction interfaces between and TapA1-EatA_NT_ and TapA2-EatA_NT_ and analysed these interactions through bacterial 2-hybrid analysis.

**Fig 3.**
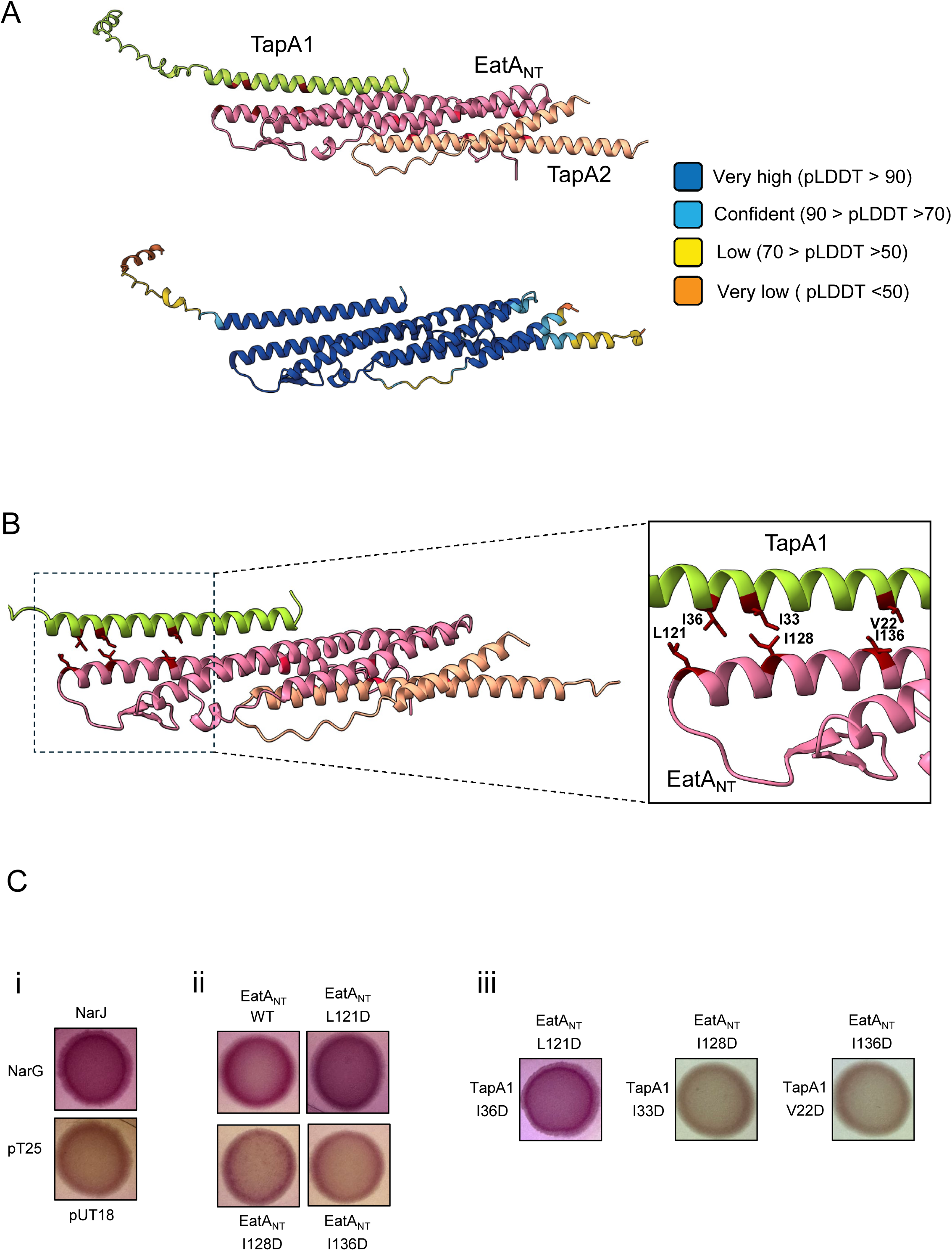
Mapping a TapA1-EatA interaction interface. (a) AlphaFold3 model of a complex containing TapA1, TapA2 and the EatA N-terminal domain (EatA_NT_: amino acids 1-180) coloured by chain (top) or pLDDT score (bottom). (b) Close-up of the TapA1-EatA_NT_ interface highlighting residues in close proximity. (c) Bacterial 2-hybrid results for (i) control proteins NarG (residues 1-42)-T18 and T25-NarJ, alongside T18 and T25 produced as non-fusion proteins; (ii) the indicated fusions of the EatA_NT_ with T18 produced alongside wild-type T25-TapA1; and (iii) the indicated point substitutions in T18- EatA_NT_ with the indicated point substitutions in T25-TapA1. All bacterial 2-hybrid analyses were carried out on the same MacConkey indicator plates, but spots have been separated for presentation.

#### The hydrophobic interface between TapA1 and EatA_NT_

The Alphafold3 prediction indicates that TapA1 interacts with the third α-helix of EatA_NT_ through a series of hydrophobic interactions. These include TapA1 V22 with EatA I136, TapA1 I33 with EatA I128. and potentially TapA1 I36 with EatA L121 (Fig 3b). To probe this interface, we first introduced individual negative charges at each of these three identified residues in EatA_NT_ which was encoded as a fusion to the N-terminus of the adenylate cyclase T18 fragment (EatA_NT_-T18), and tested the ability to interact with wild type TapA1 fused C-terminal to adenylate cyclase T25 (T25-TapA1). As seen in Fig 3c, co-production of the individual I128D and I136D substitutions of EatA_NT_ with TapA1 substantially reduced the red pigmentation on MacConkey indicator plates, consistent with a weakening of the interaction, while the L121D substitution did not. To probe this further, we next introduced individual substitutions V22D, I33D and I36D into T25-TapA, and tested them in the bacterial 2-hybrid assay against the ‘mirror’ substitutions in EatA_NT_ (Fig 3c, far-right panel). The TapA1 V22D substitution abolished the residual interaction with EatA_NT_ I136D, and likewise TapA I33D no longer showed interaction with EatA_NT_ I128D. However, the TapA1 I36D substitution still showed strong interaction with EatA_NT_ L121D, consistent with these pairs of residues being further apart in the model than the other pairings (Fig 3b). We also repeated these experiments using the reciprocal fusions, i.e. with the EatA substitutions introduced into aT25-EatA_NT_ fusion construct and TapA1 fused to T18, with essentially similar results (Fig S1a). These findings are generally consistent with the interface of TapA1 and EatA predicted by AlphaFold 3.

#### The hydrophobic interface between TapA2 and EatA_NT_

According to the model, TapA2 makes contacts with all three helices of EatA_NT_, again through a series of hydrophobic interactions including TapA2 V47 with EatA L155, TapA2 Y58 with EatA L29 and TapA2 L69 with EatA F53 (Fig 4a). To explore this further we took a similar mutagenesis/bacterial 2-hybrid strategy. We introduced single aspartate substitutions of the selected residues in the EatA_NT_-T18 fusion and first tested their interaction with wild type T25-TapA2. None of these substitutions appeared to affect the interaction as all colonies showed similar pigmentation to that seen with unsubstituted EatA_NT_-T18 (Fig 4b). We next introduced the ‘mirror’ substitutions into T25-TapA2 and repeated the analysis (Fig 4b, right hand panel). Both the TapA2 V47D - EatA_NT_ L155D and the TapA2 Y58D - EatA_NT_ L29D pairings showed negligible interaction in the 2-hybrid assay, while the TapA2 L69D substitution still showed strong interaction with EatA_NT_ F53D. Again, we repeated these experiments using the reciprocal fusions T25-EatA_NT_ and TapA2-T18, with essentially very similar results (Fig S1b). These findings are in general agreement with the interface of TapA2 and EatA predicted by AlphaFold 3, although the positioning of TapA2 L69D and EatA_NT_ F53D may show some flexibility. The findings from all the bacterial 2-hybrid analyses are summarised in Table 2.

**Fig 4.**
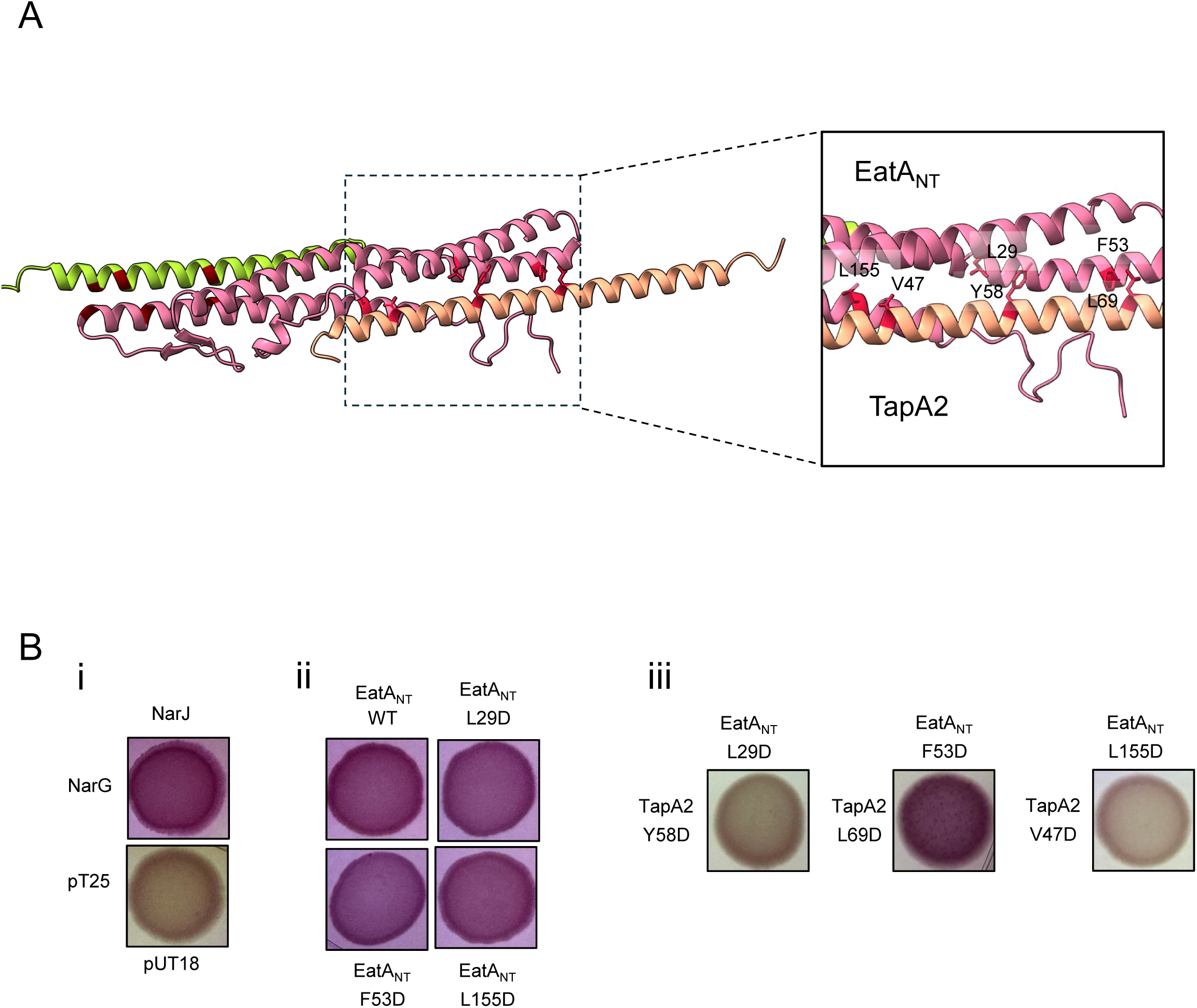
Mapping a TapA2-EatA interaction interface. (a) Close-up of the TapA2-EatA_NT_ interface highlighting residues in close proximity. (b) Bacterial 2-hybrid results for (i) control proteins NarG (residues 1-42)-T18 and T25-NarJ, alongside T18 and T25 produced as non-fusion proteins; (ii) The indicated fusions of the EatA_NT_ with T18 produced alongside wild-type T25-TapA2; and (iii) the indicated point substitutions in T18-EatA_NT_ with the indicated point substitutions in T25-TapA2. All bacterial 2- hybrid analyses were carried out on the same MacConkey indicator plates, but spots have been separated for presentation.

**Table 2.**
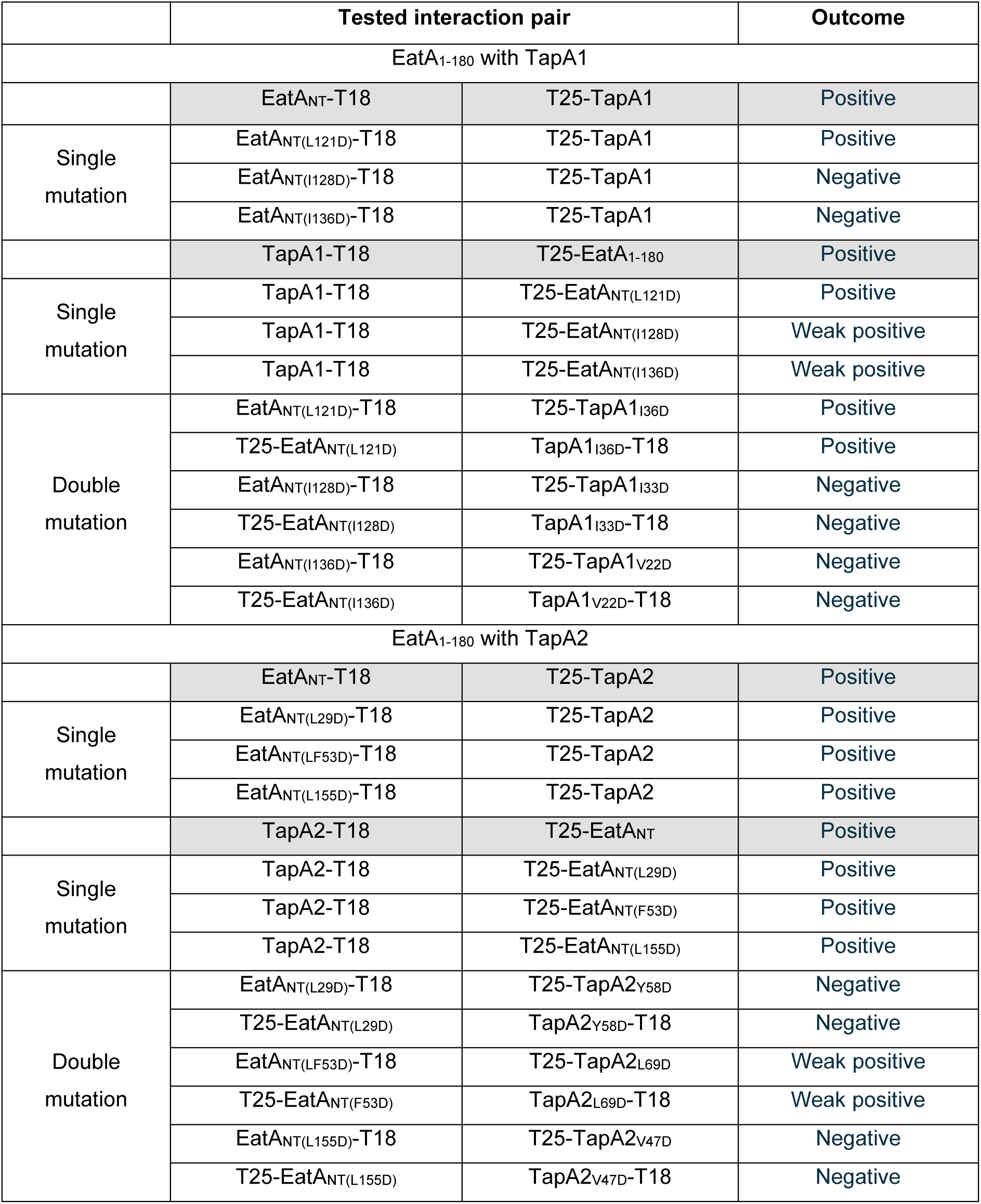
Summary of bacterial 2-hybrid interactions.

### A potential interaction site between TapA2 and EsxT?

It is clear from the literature that WXG100 family proteins may have differential functions within the T7SS. The ‘mechanistic’ WXG100 proteins, such as EsxA in the T7SSb system, or EsxAB and EsxTU in T7SSa/ESX system are necessary for operation of the secretion machinery, potentially by modulating its activity or conformation (18, 21, 35). These proteins are additionally secreted during operation of the T7SS alongside other substrate proteins (e.g. (16, 36, 37). On the other hand, other WXG100-related proteins such as the Lap proteins in the T7SSb and the TapA proteins are specific for their toxin partners but not for general operation of the secretion system (21, 26). We therefore wondered whether there could be an interaction between the EatA-TapA1-TapA2 complex and the EsxT-EsxU dimer that may occur during secretion. To explore this, we undertook AlphaFold3 modelling of the interaction between the EatA complex and EsxTU. The output of this, shown in Fig 5a, predicts a high confidence complex of all five proteins, with an interface predicted template modelling score of 0.75 and a short region of contact between EsxT and TapA2.

**Fig 5.**
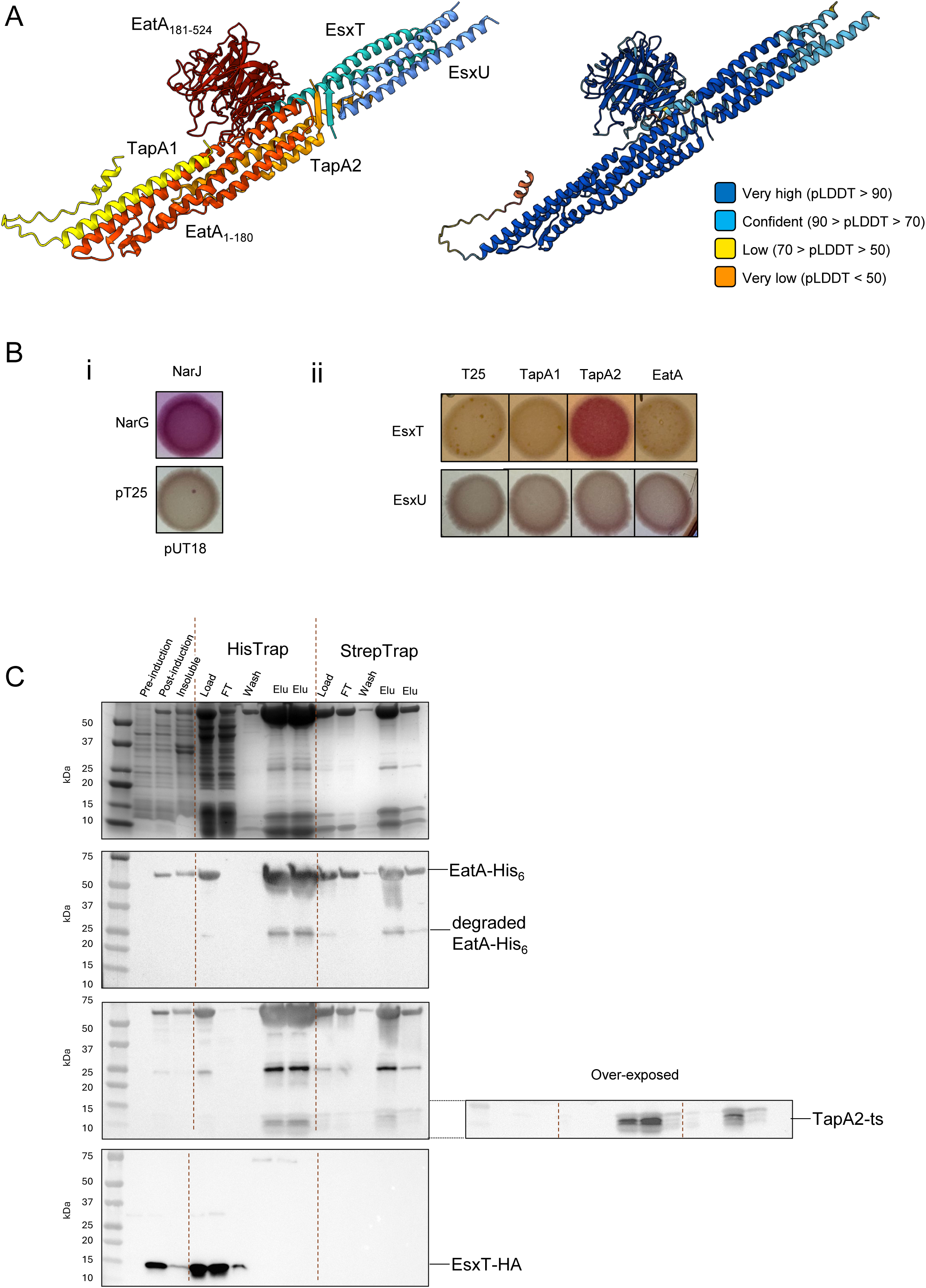
Investigating interactions between the EatA-TapA1-TapA2 complex and EsxT-EsxU. (a) AlphaFold3 model of a complex containing TapA1, TapA2, EatA, EsxT and EsxU coloured by chain (left) or pLDDT score (right). (b) Bacterial 2-hybrid results for (i) control proteins NarG (residues 1-42)- T18 and T25-NarJ, alongside T18 and T25 produced as non-fusion proteins; (ii) the indicated fusions of EsxT or EsxU with T18 (as indicated) produced alongside either T25 alone, T25-TapA1, T25-TapA2 or T25-EatA. (c) Purification of EatA-His_6_/TapA2-ts complexes. Lysates of *E. coli* co-producing EatA- His_6_, TapA2-ts, and EsxT-HA alongside untagged TapA1 and EsxU were subjected to sequential purification by HisTrap and StrepTrap chromatography. Fractions were analysed by SDS-PAGE (top panel) or western blotting with anti-His (second panel), anti-strep (third panel) or anti-HA (bottom panel) antibodies. Pre: pre-induction sample; Post: 4 hours post-induction sample; Insoluble: insoluble fraction following preparation of cell lysate, Load; sample loaded onto the column; FT: column flow-through; Wash: wash sample; Elu: elution samples from the peak fractions for each column.

To determine whether there might be a specific interaction between EsxT and TapA2, we first carried out bacterial 2-hybrid analysis, testing EsxT-T18 or EsxU-T18 fusions with each of TapA1-T25, TapA2- T25 or EatA-T25. From this, we noted that red-pigmented colonies were only visible when the EsxT fusion was present with the TapA2 fusion, consistent with the model (Fig 5b). To explore this further, we then asked whether EatA, TapA2 and EsxT could co-purify. To this end, we designed a plasmid to co-overproduce all five proteins (TapA1, TapA2, EatA, EsxT, and EsxU) in *E. coli*. After using AlphaFold3 modelling to explore where we could position tags that would not interfere with folding or potential complex formation, we introduced a C-terminal His_6_ tag on EatA, a C-terminal ts tag on TapA2 and a hemagglutinin (HA) epitope at the C-terminus of EsxT. We first isolated His_6_ tag on EatA complexes using a HisTrap column and loaded the EatA-containing fractions onto a StrepTrap column for isolation of TapA2-ts complexes. Finally, we analysed the column fractions for the presence of EatA- His_6_, TapA2-ts and EsxT-HA by western blotting (Fig 5c). While we could detect both EatA-His_6_ and TapA2-ts in the eluted fractions from both HisTrap and StrepTrap columns, consistent with them forming a complex, we did not detect any co-purifying EsxT-HA, which was only present in the flow-through fractions of the first (HisTrap) column. We conclude that if there is an interaction between EsxT and TapA2, it is not sufficiently strong to allow the proteins to co-purify.

## Discussion

In this study we have compared the interaction status of two pairs of WXG100 family proteins from *M. abscessus* – EsxT/EsxU and TapA1/TapA2. Prior results have conclusively shown that EsxT and EsxU form a heterodimer with 1:1 stoichiometry, and our co-purification experiments confirm this behaviour. By contrast we were unable to isolate heterodimers of TapA1 and TapA2, which appear only to interact indirectly through their ability to form a trimeric complex with the N-terminal domain of the EatA toxin.

At present there is no high-resolution structure for the EatA-TapA1-TapA2 complex, but there is an X-ray structure of the N-terminal domain of TelA, a T7SSb-secreted toxin, with its WXG100-like partner proteins, LapA3 and LapA4 (28). The structure reveals extensive contacts between each Lap partner and TelA, but no contact between the Lap proteins themselves. An Alphafold3 model of the EatA- TapA1-TapA2 complex likewise shows no interaction between the Tap partners (30). To test the robustness of the AlphaFold3 model we introduced single or paired negative charges into the hydrophobic interfaces between EatA_NT_ and each Tap partner and probed for interactions by bacterial 2-hybrid analysis. Our results are in general agreement with this model, and we conclude that the TapA1-TapA2-EatA_NT_ complex forms an elongated α-helical bundle similar to the T7SSb TelA complex and to other T7SSa/ESX substrates such as PE–PPE proteins (38, 39) and EspB (40).

All type VII secretion system subtypes characterised to date require a WXG100 dimer for secretion system activity. While in the T7SSb system this is a homodimer of EsxA (21, 41), in the T7SSa/ESX systems the WXG100 proteins are heterodimers. EsxT-EsxU is the heterodimeric pair associated with ESX-4, and this pair is known to be essential for secretion of at least one substrate protein (*M. tuberculosi*s CpnT; 35). We therefore wondered whether EsxT-EsxU may interact with the EatA complex which is secreted by the ESX-4 system in *M. abscessus*. Alphafold 3 modelling suggested an interaction between the EsxT-EsxU dimer and the EatA-TapA1-TapA2 trimer with high confidence, highlighting a short anti-parallel β-strand interface between TapA2 and the EsxT N-terminus. Intriguingly, bacterial 2-hybrid analysis was consistent with these two proteins interacting, however we were unable to copurify the two complexes. It should be noted that EsxA of *S. intermedius* also did not co-purify with substrate complexes (21). This might suggest that the interaction seen between TapA2 and EsxT is an artifact of the 2-hybrid approach. Alternatively, it may be that the interaction is transient and unable to survive the purification steps, or that there are components missing that may stabilise the interaction due to the heterologous expression conditions. One such component might be EccC, the central ATPase of the ESX secretion system. The EccC component of ESX-1 has been shown to bind EsxB, and since this is the only membrane component conserved across the T7SS subtypes it is generally assumed that it directly mediates secretion of all substrates (7,14). As such it is possible that the EsxT-EsxU dimer and the EatA complex are juxtaposed on the cytoplasmic ATPase domains of EccC prior to secretion. EsxT-EsxU are likely to be required for secretion of all *M. abscessus* ESX-4 substrates and might therefore be expected to make transient interactions with other complexes during secretion. Further experimental work would be required to test this hypothesis.

## Acknowledgements

We thank Drs Elisabeth Lowe and Patrick Moynihan for helpful discussions.

## Author contributions

Conceptualization (TP, EKEL, ERB); Investigation (EKEL, KB); Writing (TP, EKEL); Supervision (TP, ERB) and Funding acquisition (TP).

