## supplementary Table and Figure for "Characterizing the interaction of a type VII-secreted antimycobacterial toxin with its small helical partner proteins"

**SUPPLEMENTARY INFORMATION**

**Table S1. Oligonucleotide primers used in this study**

| Name | Sequence (5' - 3') | Usage / Construct |
|---|---|---|
| T18_Fwd | GAGCGGATAACAATTTCAC | Sequencing of pUT18 constructs, flank the multiple cloning site. |
| T18_Rev | CTGGCGACGCGCCTCGGTGCC | |
| T25_Fwd | GGATGTGCTGCAAGGC | Sequencing of pT25 constructs, flank the multiple cloning site. |
| T25_Rev | CCGCATCTGTCCAACTTCC | |
| QE70_Fwd | CCCGAAAAGTGCCACCTG | Sequencing of pQE70 constructs, flank the MCS. |
| QE70_Rev | GTTCTGAGGTCATTACTGGAT | |
| T18-esxU_F | TGACCATGATTACGCCAAGCGCTGTTTTTCAGAATGACCTG | pUT18-*esxU* |
| T18-esxU_R | GTCGACCTGCAGGCATGCAAG AAGGTCTGGCTCTCGTC | |
| T18-esxT_F | TGACCATGATTACCCAAGCAGCCAGATTACTTACAACCACGCGAGATTGACG | pUT18-*esxT* |
| T18-esxT_R | GTCGACCTGCAGGCATGCAAGTGGTGCCAGGCGCCGGC | |
| V22D_for | GATATGGATATTGAGGCGGCGGGA | pUT18-*tapA1*$_{V22D}$ |
| V22D_rev | ACTACGATCTGCGACACTACGAATC | pT25-*tapA1*$_{V22D}$ |
| I33D_for | GATGCCGAGATTGATCGGCGTG | pUT18-*tapA1*$_{I33D}$ |
| I33D_rev | GGCTTCGCGTCCGGCCGCC | pT25-*tapA1*$_{I33D}$ |
| I136D_for | GATCTGCACACCCGGGCGCAGGA | pUT18-*eatA*$_{1\text{-}180(I136D)}$ |
| I136D_rev | CCCCTGCAGCCACCTCCGAC | pT25-*eatA*$_{1\text{-}180(I136D)}$ |
| I128_for | GATCGTCGGAGGTGGCTGCAGGG | pUT18-*eatA*$_{1\text{-}180(I128D)}$ |
| I128_rev | TGCCCTCTTCAGACGGTCCAAG | pT25-*eatA*$_{1\text{-}180(I128D)}$ |
| L121_for | GATGACCGTCTGAAGAGGGCAATAC | pUT18-*eatA*$_{1\text{-}180(L121D)}$ |
| L121_rev | GCCAGAATGCGATCGTCGCTC | pT25-*eatA*$_{1\text{-}180(L121D)}$ |
| L155_for | GATCATGAAATCGCCAAGATTTAAGC | pUT18-*eatA*$_{1\text{-}180(L155D)}$ |
| L155_rev | CTTCCGAGCGATCGCGTTAT | pT25-*eatA*$_{1\text{-}180(L155D)}$ |
| L29_for | GATGACACCGCAGTCAACGGTGC | pUT18-*eatA*$_{1\text{-}180(L29D)}$ |
| L29_rev | CTGCG TAGCTTGGTCCCCCC | pT25-*eatA*$_{1\text{-}180(L29D)}$ |
| F53_for | GATCGGACGCGGCTGAAAAGCTATATC | pUT18-*eatA*$_{1\text{-}180(F53D)}$ |
| F53_rev | CGCCTCGACAAACTGGCCGG | pT25-*eatA*$_{1\text{-}180(F53D)}$ |
| V47D_for | GATTCGGCATCCGGTGACCACGT | pT25-t*apA2*$_{V47D}$ |
| V47D_rev | GGCGGACACAGCCGCAACGG | |
| Y58D_for | GATGGTGCGCTTCTTCAGGAGGC | pT25-t*apA2*$_{Y58D}$ |
| Y58D_rev | TTCCTTGGCGACGTGGTCACC | |

| | | |
|---|---|---|
| L69D_for | GATGGCCGCGCGG TT CAAGCTTT | pT25-t*apA2*$_{\text{L69D}}$ |
| L69D_rev | GAACGCTGCGGCCTCCTGAAG | |
| L56D_rev | GGGCTGCCTCACCTGGGCCT | |
| I36D_for | GATGATCGGCGTGCGACGGCA | pUT18-t*apA1*$_{\text{I36D}}$ |
| I36D_rev | CTCGGCGATGGCTTCGCGTC | pT25-t*apA1*$_{\text{I36D}}$ |
| Bbackbone_R | TGTAGCTCCTGTTCATCACTCATGCTTAATTTCTCCTCTTTAATGAATTCTGTGT | |
| B1_F | AAGAGGAGAAATTAAGCATGAGTGATGAACAGGAGCTACAAC | |
| B1_R | TGTGGATGACTCCAAGCACTCATATTCTGTGGGCCGGAGG | |
| plasmidE_for | CATCACCATCACCATCACTAAGCTTAATTAGC | Used Plasmid B as template; KLD; pQE70-*tapA1-tapA2-eatA*$_{\text{1-180}}$ (Plasmid E) |
| plasmidE_rev | TCGATGGCCACTGAAGTCCG | |
| plasmidE2.0_for | TTAGTGATGGTGATGGTGATGTCGATG | pQE70-*tapA1-tapA2-eatA*$_{\text{1-180}}$-*TAA* (Plasmid E2.0) |
| plasmidE2.0_rev | TAAGCTTAATTAGCTGAGCTTGGACTCCTGTTGAT | |
| PQE70_for | TAAGCTTAATTAGCTGAGC | pQE70-*tapA1-tapA2* |
| PQE70_rev | GCTTAATTTCTCCTCTTTAATG | |
| wxg_for | TTAAAGAGGAGAAATTAAGCATGAGTGATGAACAGGAGC | |
| wxg_rev | TTAGTGATGGTGATGGTGATGGCGGAACCTCTGGAATTG | |
| rbspluswxg2ts_F | CATCACCATCACCATCACTAATGGAGAGGGGCAACCAATG | |
| rbspluswxg2ts_R | AGCTCAGCTAATTAAGCTTACTTCTCAAATTGTGGATGAGACC | |
| pqe70_for | ATCCACAATTTGAGAAGTAGGCTTAATTAGCTGAGCTTG | pQE70-*esxU-esxT* |
| pqe70_rev | CATGCTT AATTTCTCCTCTTTAATG | |
| nostartEsxUnostop_F | AAGAGGAGAAATTAAGCATGGCTGTTTTTCAGAATGACCTG | |
| nostartEsxUnostop_R | CATGGTTAATTTCTCCTCTTTAATTTATTAGTGATGGTGATGGTGATGGAAGGTCTGGCTCTCGTC | |
| nostartEsxTnostop_F | CATCACCATCACCATCACTAATAAATTAAAGAGGAGAAATTAACCATGAGCCAGATTACTTACAACCACGGCGAGATTGACG | |
| nostartEsxTnostop_R | TGTGGATGACTCCAAGCACTGTGGTGCCAGGCGCCGGC | |
| twinstrep_F | AGTGCTTGGAGTCATCCAC | |

| | | |
|---|---|---|
| twinstrep_R | CTACTTCTCAAATTGTGGATGAG | |
| Bbackbone_F | TCGAAGTAAACCCACATCAACATCACCATCACCATCACTAAGCT | pQE70-*tapA1-tapA2-eatA* (Plasmid B) |
| Bbackbone_R | TGTAGCTCCTGTTCATCACTCATGCTTAATTTCTCCTCTTTAATGAATTCTGTGT | |
| B1_F | AAGAGGAGAAATTAAGCATGAGTGATGAACAGGAGCTACAAC | |
| B1_R | TGTGGATGACTCCAAGCACTCATATTCTGTGGGCCGGAGG | |
| B2_F | CCTCCGGCCCACAGAATATGAGTGCTTGGAGTCATCCACAATTC | |
| B2_R | TACATATTCTGTGGGCCGGACTACTTCTCAAATTGTGGATGAGACCAG | |
| B3_F | ATCCACAATTTGAGAAGTAGTCCGGCCCACAGAATATGTAGGC | |
| B3_R | TAGTGATGGTGATGGTGATGTTGATGTGGGTTTACTTCGATCAATTGGG | |
| 0404-plasmidB_F | TACCCATACGATGTTCCAGATTACGCTTAAGCTTAATTAGCTGAGCTTGGACTCC | pQE70-*tapA1-tapA2-eatA-esxU-esxT* |
| 0404-plasmidB_R | TTTCAGTTCCTCCCTTGTCCCCAGGCCCGGCTTAGTGATGGTGATGGTGATGTTGATG | |
| EsxUTfrgDNA_F | GCCGGGCCTGGGGACAAGGGAGGAACTGAAATGGCTGTTTTTCAGAATGACCTGGCGTTGC | |
| EsxUTfrgDNA_R | TTAAGCGTAATCTGGAACATCGTATGGGTAGTGGTGCCAGGCGCCGGC | |

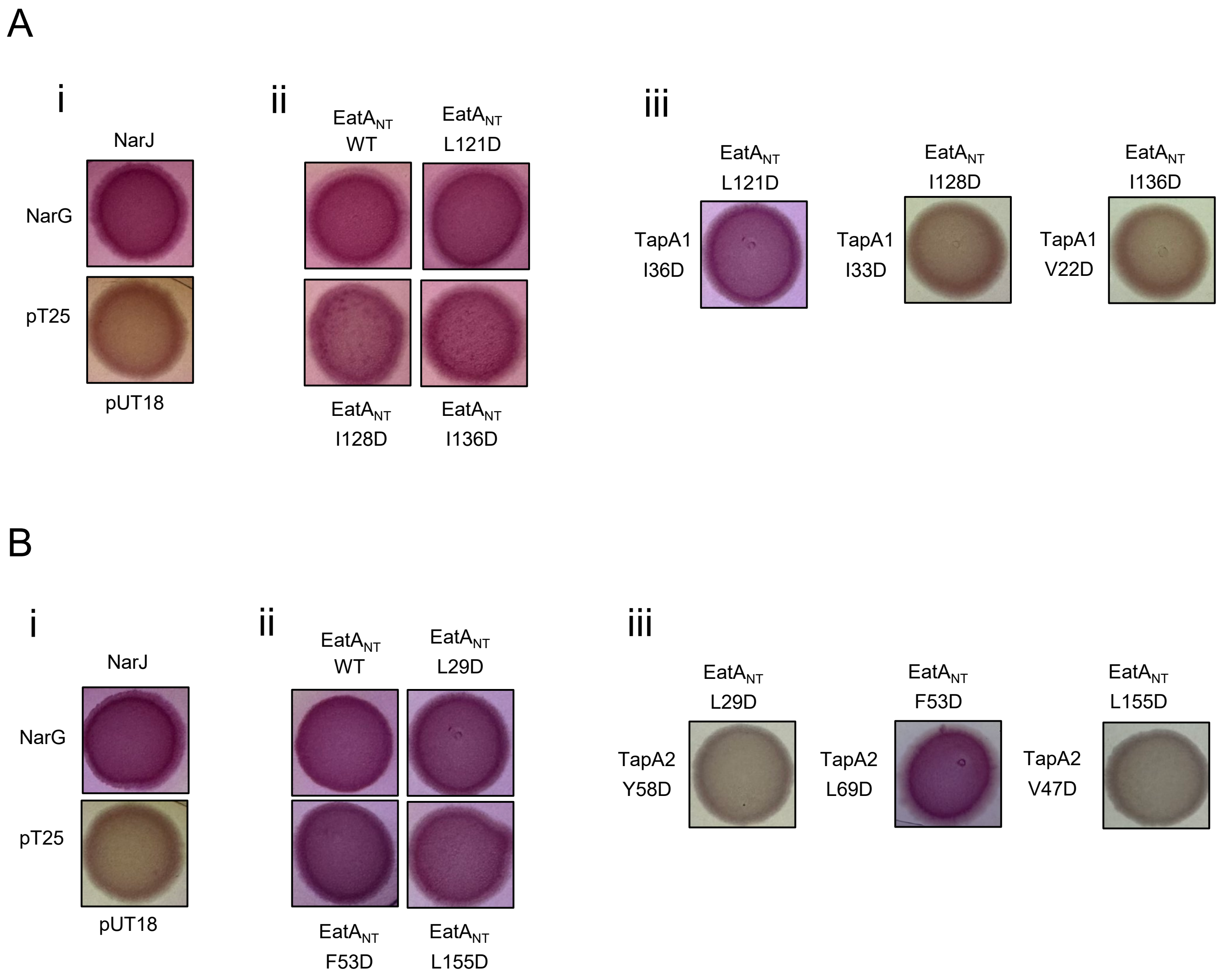


Fig S1. Bacterial 2-hybrid results (a) for (i) control proteins NarG (residues 1-42)-T18 and T25-NarJ, alongside T18 and T25 produced as non-fusion proteins; (ii) The indicated fusions of the EatANT with T25 produced alongside wild-type TapA1-T18; and (iii) the indicated point substitutions in EatANT-T25 with the indicated point substitutions in TapA1-T18. Note that the controls in (i) are the same as for Fig 3C as all of the analyses shown in Fig 3c and Fig S1c were carried out on the same MacConkey indicator plates, and (b) (i) control proteins NarG (residues 1-42)-T18 and T25-NarJ, alongside T18 and T25 produced as non-fusion proteins; (ii) the indicated fusions of the EatANT with T25 produced alongside wild-type TapA2-T18; and (iii) the indicated point substitutions in EatANT-T25 with the indicated point substitutions in TapA2-T18. Note that the controls in (i) are the same as for Fig 4b as all of the analyses shown in Fig 4b and Fig S1b were carried out on the same MacConkey indicator plates.
